# MEK/mTOR-dependent D1 dopamine receptor activation induces local protein synthesis via eEF2 dephosphorylation in neurons

**DOI:** 10.1101/447714

**Authors:** Orit David, Iliana Barrera, Bella Koltun, Sivan Ironi, Shunit Gal-Ben-Ari, Kobi Rosenblum

**Affiliations:** Sagol Department of Neurobiology, University of Haifa, Haifa 3498838, Israel; Center for Gene Manipulation in the Brain, University of Haifa, Haifa 3498838, Israel

**Author notes:** O.D. and I.B. contributed equally to this work. Author to whom correspondence should be addressed: Kobi Rosenblum, PhD Sagol Department of Neurobiology Center for Gene Manipulation in the Brain University of Haifa. Haifa 3498838, Israel.

## Abstract

Neuromodulators in general, and dopamine in particular, define brain and neuronal states in different ways including regulation of global and local mRNA translation. Yet, the signaling pathways underlying the effects of dopamine on mRNA translation are not clear. Here, using genetic, pharmacologic, biochemical, and imaging methods, we tested the hypothesis that dopamine regulates phosphorylation of the eukaryotic elongation factor 2 (eEF2). We found that activation of dopamine receptor D1 but not D2 leads to rapid dephosphorylation of eEF2 at Thr^56^ in cortical primary neuronal culture and *in vivo* in a time-dependent manner. Additionally, NMDA receptor, mTOR, and ERK pathways are upstream to the D1 receptor-dependent eEF2 dephosphorylation and essential for it. Furthermore, D1 receptor activation resulted in a major reduction in dendritic eEF2 phosphorylation levels together with a correlative increase in local mRNA translation. These results reveal the role of eEF2 in dopamine regulation of local mRNA translation in neurons.

**One-sentence summary:** D1 receptor activation increases protein synthesis in dendrites by inactivating eEF2K in an ERK2/mTOR-dependent manner.

## Introduction

Dopamine is an important neuromodulator involved in motivation, addiction, and learning and memory, which modulates neuronal activity and synaptic plasticity in different brain areas (Chafee and Goldman-Rakic, 1998, Waelti et al., 2001). It exerts its complex effect via regulation of cellular and molecular networks. Modulation of long-term memory by dopamine requires *de novo* local protein synthesis (Schicknick et al., 2008), a process controlled by translation factors.

The translation factor elongation factor 2 (eEF2) and its kinase (eEF2K)play a central role in the elongation phase of mRNA translation. eEF2K phosphorylates eEF2 on Thr^56^ and thereby inactivates it, leading to reduction in the rate of mRNA translation. Since eEF2K activity is regulated by Ca^2+^/calmodulin, elevation in intracellular calcium by synaptic activation receptors such as NMDA receptor or G-coupled receptors (e.g. metabotropic glutamate receptors) results in induced neuronal activity-dependent phosphorylation of eEF2 (Proud, 2015, Barrera et al., 2008, Heise et al., 2017, Im et al., 2009, Sutton et al., 2007). Furthermore, general translation is reduced in dendrites due to eEF2K activity, while certain synaptic proteins are selectively translated at the synapse (Heise et al., 2014).

eEF2K can be deactivated by other kinases such as p70S6 kinase (S6K) or p90 ribosomal kinase (p90 RSK), which are activated in response to changes in mammalian target of rapamycin-signaling (mTOR) or extracellular-regulated kinase (ERK) signaling respectively. These kinases inactivate eEF2K by phosphorylation on its Ser366 residue (Browne et al., 2004, Redpath et al., 1993, Wang et al., 2001).

Regulation of protein synthesis by neurotransmitters, especially at the phase of elongation, has been shown as a mechanism to explain the effects of antidepressants. For example, ketamine, a NMDA receptor non-competitive antagonist, has a fast, potent, and long-lasting antidepressant effect mediated by the reduction in eEF2 phosphorylation via eEF2K (Adaikkan et al., 2018, Duman and Voleti, 2012). Interestingly, drugs targeting the function of dopamine receptors and their interaction with NMDA receptors have become an important field of research to develop better antidepressant medications (Miller et al., 2014, Raab-Graham and Niere, 2017). However, the mechanism by which dopamine receptors regulate mRNA translation during the elongation phase remains poorly understood.

In this work, we found that D1 but not D2 receptor activation increases local protein synthesis in dendrites by eEF2 dephosphorylation. This increase in local protein synthesis in cortical neurons is mediated by the D1 receptor and requires NMDA receptor-dependent activation of the MEK/mTOR signaling pathways that lead to inactivation of eEF2K by phosphorylation on Ser^366^ residue resulting in eEF2 dephosphorylation. Furthermore, using eEF2K knockout mice, we show that eEF2K/eEF2 is the main pathway for D1-dependent increase in local protein synthesis.

## Results

### D1 receptor activation induces eEF2 dephosphorylation in vitro and in vivo

The mTOR and MEK-ERK pathways are regulators of eEF2K (Browne et al., 2004, Proud, 2015, Redpath et al., 1993, Wang et al., 2001). Since activation of dopamine receptors can modulate both pathways in various types of cells and different behaviors (Gangarossa et al., 2012, Nagai et al., 2007, Schicknick et al., 2008, Schicknick et al., 2012, Biever et al., 2015), we asked whether stimulation of the different dopamine receptors could induce changes in eEF2K and its sole downstream known target, eEF2 (Kenney et al., 2015). Using primary cortical neuronal cultures, we found that treatment with D1 receptor agonist SKF38393, but not the D2 agonist, quinpirole, induced rapid dephosphorylation of eEF2 which lasted for one hour. Pretreatment of the cells with SCH23390, a D1 receptor antagonist, blocked the D1 receptor-induced dephosphorylation of eEF2 (Figure 1A), while pretreatment with D2 receptor antagonist, eticlopride, did not have any effect (Figure 1B). As expected, SKF38393 treatment of primary cultures induced ERK1/2 phosphorylation as reported before (Gangarossa G and Valjent E., 2012; David et al 2014) (Figure1 supplement 1). Moreover, ERK1/2 activation correlated with dephosphorylation of eEF2 by D1 receptor stimulation (r= -0.50, p<0.05, Pearson’s correlation) (Figure1 supplement 1).

**Figure 1.**
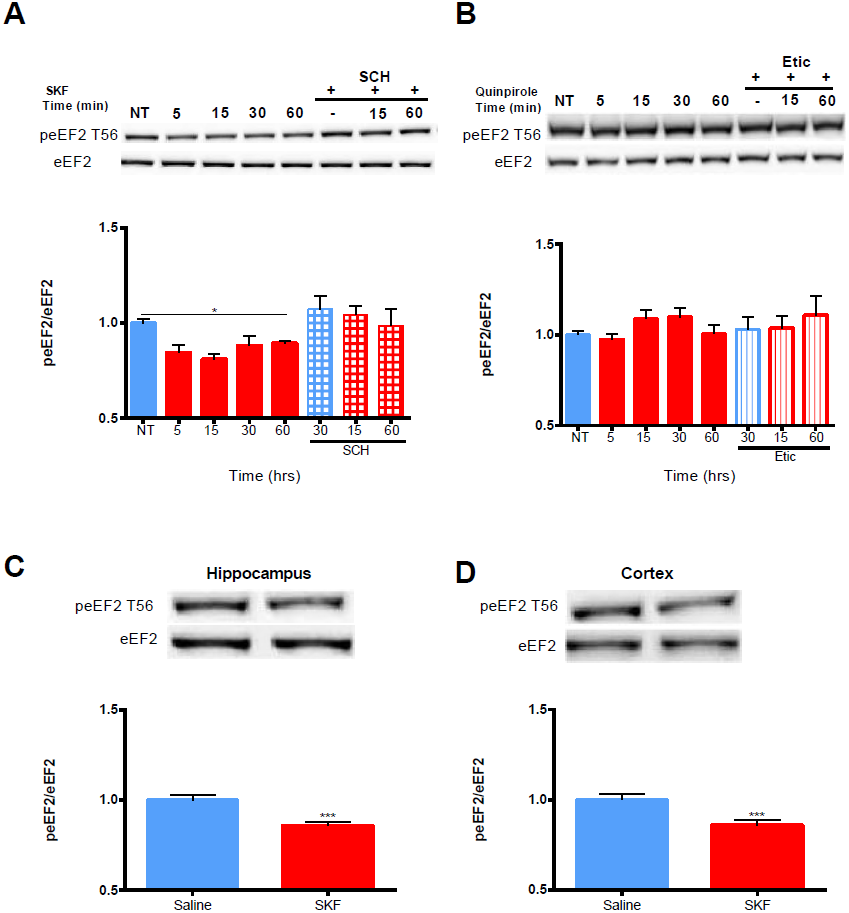
D1 but not D2 receptor activation reduces eEF2 phosphorylation in cortical neurons and in vivo. **(A)** Cortical primary cultures were treated with D1 receptor antagonist SKF38393 (25μM) for the indicated time periods with or without pre-incubation with D1 receptor antagonist SCH23390 (10μM, 30 min) followed by incubation with SKF38393. **(B)** Cortical primary cultures were treated as in (A) with D2 agonist quinpirole (10μM) for the indicated time periods with or without preincubation with D2 antagonist eticlopride (20μM, 30 min). Data are means ± SEM of five independent cultures. **(C and D)** Quantification of the ratio of phosphorylated to total eEF2 in hippocampal and cortical extracts from C57BL/6 male mice injected i.p. with SKF38393 (5mg/Kg) and sacrificed 15 minutes later. Representative immunoblots are shown. Data are means ± SEM of 12-19 mice per condition, ^*^p<0.05, ^***^p<0.001. See also Figure 1-source data 1 and Figure 1-figure supplement 1-3. SKF, SKF38393; SCH, SCH23390; etic, eticlopride. **Source data 1-Statistical analysis of Figure 1** **Figure 1A.** One-way ANOVA (F_(7,44)_=5.054, p<0.0001); Post-hoc test compared with control: SKF5min, p=0.04; SKF15min, p=0.026; SKF60, p=0.04; SKF15min compared to SKF15+SCH, p=0.015; SKF60, compared to SKF60+SCH, p=0.04. **Figure 1B.** One-way ANOVA (F_(7,37)_=0.847, p=0.556. **Figure 1C.** Independent samples t-test: saline, n = 12 mice; SKF38393, n=12 mice; T _(22)_ =3.81, p=0.001. **Figure 1D.** Independent samples t-test: saline, n = 12 mice; SKF38393, n=12 mice; T _(29)_ =3.33, p=0.002.

Following the results we obtained in primary cultures, we further examined whether D1 activation induces dephosphorylation of eEF2 *in-vivo* as well. Western blot analyses of both cortex and hippocampus extracts from SKF38393-injected C57BL/6J mice (i.p. injection) showed reduction in eEF2 phosphorylation after 15 minutes (Figure 1C and figure 1D), similar to the data obtained in primary cultures (Figure 1A). In agreement with previous results, ERK2 activation was increased in both hippocampus and cortex of these SKF38393-treated mice compared to controls (Figure 1 supplement 1 C and Figure 1 supplement 1 D). These results suggest that dopamine regulates eEF2 phosphorylation both in vivo and in vitro via D1 but not D2 receptor.

### eEF2 dephosphorylation by D1 receptor activation is dependent on the NMDA receptor

Following the correlation between ERK2 activation as indicated by its phosphorylation state and eEF2 dephosphorylation (Figure 1 supplement 1 A and Figure 1 supplement 1B), and since ERK is a coincidence detector of dopamine and the NMDA receptor (Kaphzan 2006, David 2014), we further asked whether the NMDA receptor has a role in D1 receptor-dependent dephosphorylation of eEF2. To test this, primary cortical neuronal cultures were pretreated for 30 min with APV, a NMDA antagonist, followed by 15 or 60 min treatment of D1 receptor agonist SKF38393. Treatment of the cells with APV prevented the dephosphorylation of eEF2 following SKF38393 treatment, both after 15 and 60 min (Figure 2A).

**Figure 2.**
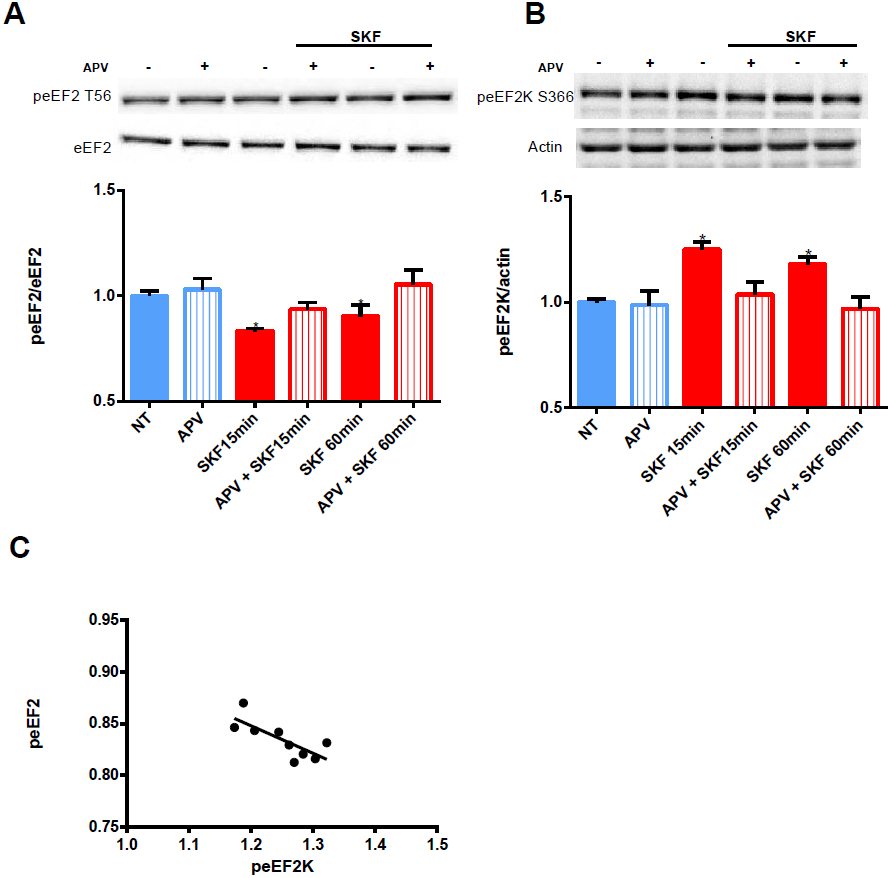
NMDA receptor mediates D1 receptor-dependent eEF2 dephosphorylation. Representative blots and quantification of the ratios of **(A)** phosphorylated to total eEF2 and **(B)** phosphorylated eEF2K to β-actin from cortical primary cultures incubated with SKF38393 (25μM) for 15 or 60 min, with or without 30 min pretreatment with NMDA receptor antagonist APV (40μM). **(C)** Negative correlation between peEF2Thr^56^ and peEF2KSer^366^ in cortical cultures treated with SKF38393 for 15 min. Data are means ± SEM of six independent cultures.^*^p<0.05. SKF, SKF38393. **Source data 1. Statistical analysis of Figure 2** **Figure 2A.** One way ANOVA F_(538)_=2.631, p=0.03. Post-hoc test compared with control, SKF 15min, p=0.04; SKF 60min, p=0.04; post hoc for SKF 15min, compared to SKF 15 min+APV, p=0.04. **Figure 2B.** One way ANOVA F_(538)_=2.669, p=0.02. Post-hoc test compared with control, SKF 15min, p=0.03; SKF 60min, p=0.04; post-hoc for SKF 15min compared to SKF 15 min+APV, p=0.04; post-hoc for SKF 60min compared to SKF 60min+APV, p=0.04. **Figure 2C.** Pearson’s correlation, r=-0.76, p<0.05.

Since eEF2K inhibits eEF2 activity, and is negatively regulated by phosphorylation on Ser^366^, we further asked whether treatment with SKF38393 increases eEF2K phosphorylation on Ser^366^ at the same time points when we observed eEF2 dephosphorylation. Indeed, treatment of primary cultures with SKF38393 clearly induced phosphorylation of eEF2K on Ser^366^, which was blocked by pre-incubation with NMDA receptor antagonist, APV (Figure 2B). As expected, APV also abolished ERK1/2 activation as shown in previous studies (Kaphzan et al., 2006; David et al., 2014) (Figure 2 supplement 1A). Furthermore, eEF2 dephosphorylation showed negative correlation with eEF2K phosphorylation (r=0.76, p<0.05, Pearson’s correlation) (Figure 2C). These results suggest that NMDA receptor activation plays a role in D1 receptor-dependent regulation of eEF2 and eEF2K activities.

### Dopamine D1 receptor activation induces eEF2 dephosphorylation in an ERK-and mTOR-dependent manner

Since both MEK-ERK and mTOR can lead to cross-activation and pathway convergence on substrates, as in the case of eEF2K phosphorylation (Redpath and Proud, 1993a; Wang et al., 2001; Browne et al., 2004; Browne and Proud, 2004; Lenz and Avruch, 2005), we sought to differentiate between these pathways, and test whether one of them is more dominant in the inhibition of eEF2K and inducing D1 receptor-dependent eEF2 dephosphorylation in neurons. Pre-incubation of cortical neurons with U0126 compound, a MEK inhibitor, blocked the phosphorylation of eEF2K at Ser^366^, inhibited eEF2 dephosphorylation (Figure 3A), and reduced dopamine D1 receptor-dependent activation of ERK1/2 and the phosphorylation of S6K at Thr^389^ (Figure 3C). Interestingly, incubation of the cells with rapamycin alone resulted in enhancement of eEF2 phosphorylation and reduction in S6K phosphorylation at the Thr^389^ residue (Figure 4A and Figure 4B) without affecting ERK1/2 phosphorylation (Figure 4B). Pre-incubation with U0126 markedly reduced the phosphorylation of ERK1/2 and S6K (Figure 3B). These findings imply that the MEK-ERK pathway is upstream to the mTOR pathway following D1 receptor stimulation.

**Figure 3.**
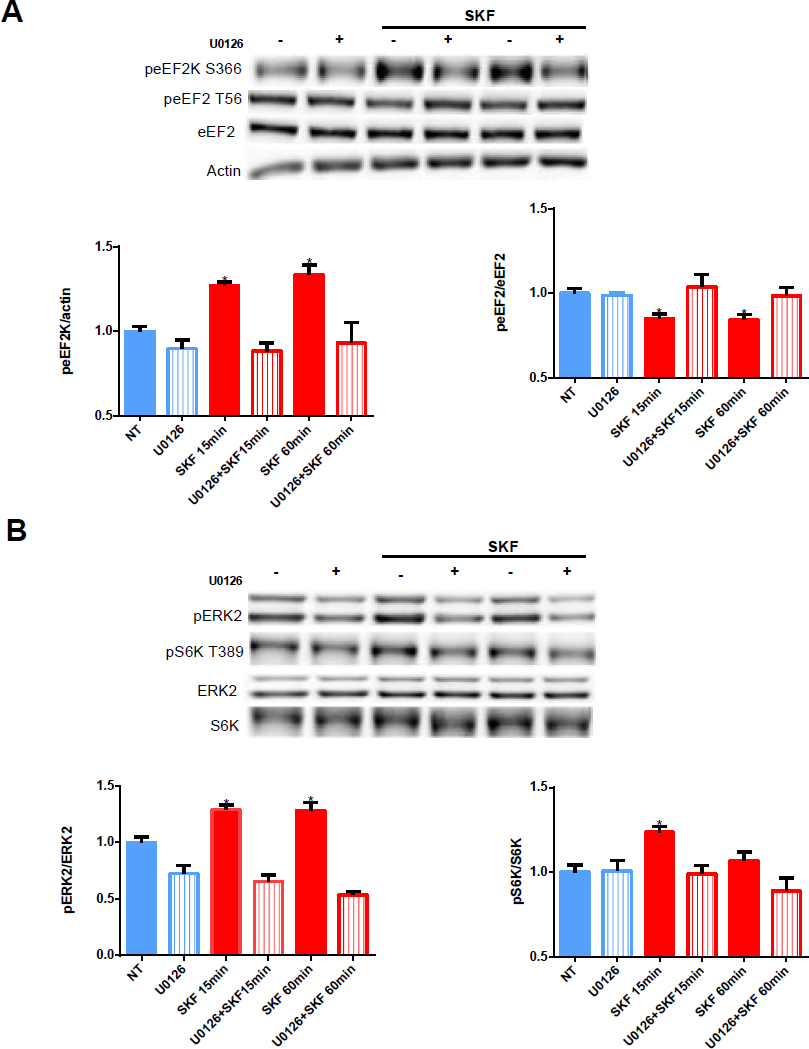
Inhibition of MEK/ERK pathway increases eEF2K activity and reduces S6K phosphorylation. **(A)**Representative blots and quantification of the ratios of phosphorylated to total eEF2 and phosphorylated eEF2K to β-actin from cortical primary cultures incubated with SKF38393 (25μM), for 15 or 60 min with or without pretreatment with MEK inhibitor U0126 (20μM) for 30 min. **(B)** Representative blots and quantification of phosphorylated to total ERK1/2 and phosphorylated S6K from cortical primary cultures treated as in A. Data are means ± SEM of five independent cultures, ^*^p<0.05. See also Figure 3-source data 1 and Figure 3-figure supplement 1. **Source data 1. Statistical analysis of Figure 3** **Figure 3A.** For eEF2K: One way ANOVA F_(5,22)_=8.624, p<0.0001. Post-hoc test compared to Control; SKF15min, p=0.03; SKF60min, p=0.012; post-hoc for SKF15min compared to SKF15+U0126, p=0.04; post-hoc for SKF60min compared to SKF60+U0126, p=0.006. For peEF2: One way ANOVA F_(5,22)_=4.506, p=0.006. Post-hoc test compared to Control; SKF15min, p=0.04; SKF60min, p=0.03; post-hoc for SKF15min compared to SKF15+U0126, p=0.04; post-hoc for SKF60min, compare to SKF60min+U0126, p=0.03. **Figure 3B.** For ERK: One way ANOVA F_(5,22)_=21.463, p<0.0001. Post-hoc test compared to Control; U0126, p=0.02; SKF15min, p=0.04; SKF60min, p=0.03; post-hoc for SKF15min compared to SKF15+U0126, p<0.0001; post-hoc for SKF60min, compared to SKF60+U0126, p<0.0001. For pS6K: One way ANOVA F_(5,22)_=4.179, p=0.008. Post-hoc test compared to Control; SKF15min, p=0.02; post-hoc for SKF15min, compared to SKF15+U0126, p=0.04.

**Figure 4.**
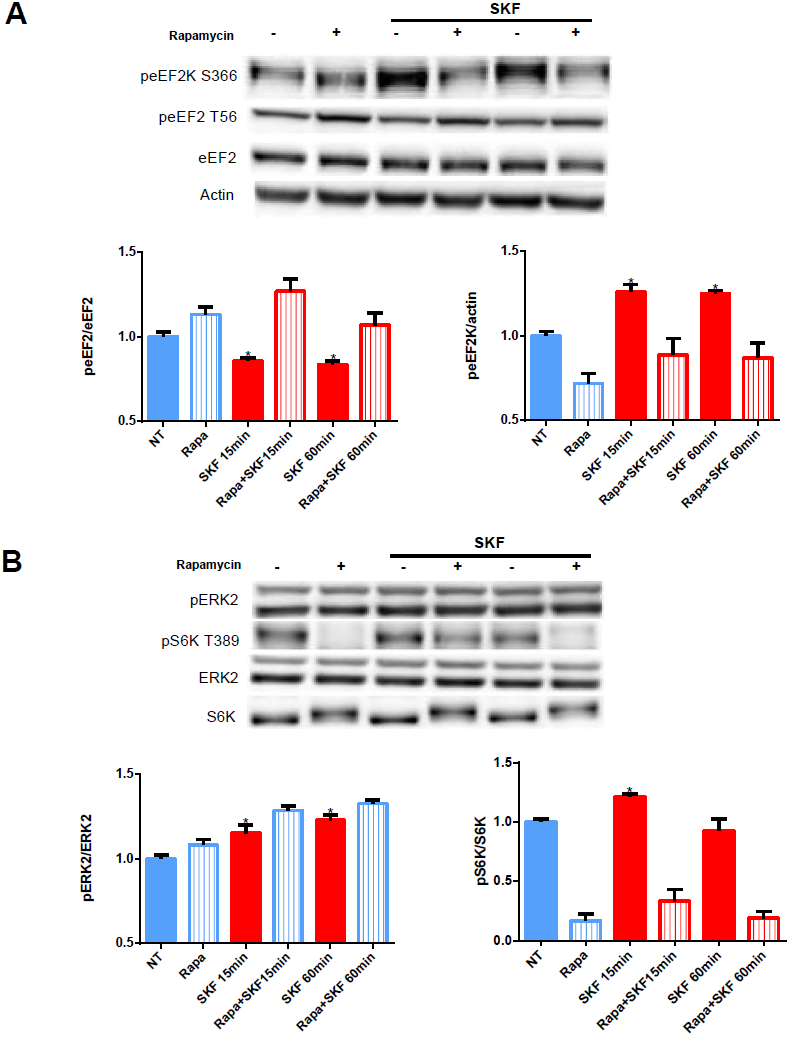
D1 receptor-dependent dephosphorylation of eEF2 requires mTOR pathway activation. (A) Representative blots and quantification of the ratios of phosphorylated to total eEF2 and phosphorylated eEF2K to β-actin from cortical primary cultures incubated with SKF38393 (25μM) for 15 or 60 min with or without pretreatment with mTOR inhibitor rapamycin (100nM) for 30 min. **(B)** Representative blots and quantification of phosphorylated to total ERK2 and phosphorylated S6K from cortical primary culture treated as in A. Data are means ± SEM of six independent cultures, ^*^p<0.05. See also Figure 4-source data 1 and Figure 4-figure supplement 1-3. **Source data 1. Statistical analysis of Figure 4** **Figure 4A.** For eEF2: One way ANOVA F_(5,39)_=9.297, p<0.0001. Post-hoc test compared to Control; SKF15min, p=0.03; SKF15+Rapa, p=0.02; SKF60min, p=0.03; post-hoc for SKF15min compared to SKF15+Rapa, p<0.0001; post-hoc for SKF60min compared to SKF60+Rapa, p=0.04. For eEF2K: One way ANOVA F_(5,39)_=6.573, p<0.0001. Post-hoc test compared to Control; Rapa, p=0.04; SKF15min, p=0.02; SKF60min, p=0.03; post-hoc for SKF15min, compared to SKF15+Rapa, p=0.04; post-hoc for SKF60min, compared to SKF60+Rapa, p=0.04. **Figure 4B.** For ERK: One way ANOVA F_(5,39)_=4.833, p=0.002. Post-hoc test compared to Control; SKF15min, p=0.02; SKF60min, p=0.005; SKF60+Rapa, p=0.036. For pS6K: One way ANOVA F_(5,39)_=39.56, p<0.0001. Post-hoc test compared to Control; Rapa, p<0.0001; SKF15min, p=0.02; post-hoc for SKF15min compared to SKF15+Rapa, p<0.0001; post-hoc for SKF60min compared to SKF60+Rapa, p<0.0001.

### Dopamine D1 receptor dephosphorylates eEF2^Thr56^ predominantly in dendrites

Regulation of eEF2K activity and reduction of eEF2 phosphorylation by synaptic receptors such as the NMDA receptor serves as one possible way of mRNA translation in neuronal dendrites (Autry et al., 2011). To examine whether dopaminergic transmission can also regulate dendritic eEF2 phosphorylation, we performed immunocytochemical analysis of cortical neurons from wild-type mice treated with SKF38393 for 15 minutes. The results revealed a reduction in phospho-eEF2 immunoreactivity in most of the neurons analyzed. The reduction was measured in both neuronal soma (Figure. 5A) and dendrites (Figure. 5B), with a more pronounced effect in the dendrites. Phospho-eEF2 antibody specificity was tested in primary cultures derived from eEF2K-KO mice. No immunoreactivity was detected in cortical neurons from eEF2K-KO cultures (Figure 5 supplement 1). These results show that D1 receptor activation reduces eEF2 phosphorylation in dendrites, suggesting its potential role in local protein synthesis.

**Figure 5.**
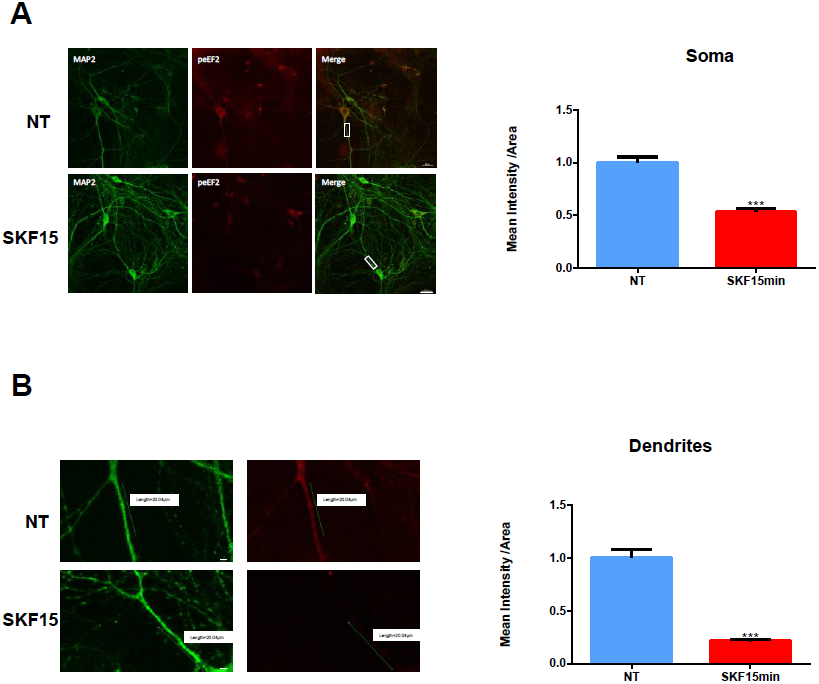
D1 receptor activation induces eEF2 dephosphorylation in dendrites more than in cell soma. **(A)** Immunofluorescence and quantification of phospho- eEF2 (Thr^56^) (red). Mean intensity in soma of MAP2 (green) positive cells from primary cortical cultures treated with SKF38393 (25μM) for 15 min. Scale bar, 20μm. Images represent 15 to 20 neurons from four independent cultures. **(B)** Immunofluorescence and quantification of phospho-eEF2 (red). Mean intensity in dendrites of MAP2 (green) positive cells from primary cortical cultures treated with SKF38393 (25μM) for 15 min. Scale bar, 10μm. Images represent 15 to 20 neurons from four independent cultures. Data are means ± SEM; ^***^p<0.0001. See also Figure 5-source data 1 and Figure 5 supplement 1. **Source data 1. Statistical analysis of Figure 5.** **Figure 5A.** Mann-Whitney test, U = 8839, Z = -8.994, P < 0.0001. **Figure 5B.** Mann-Whitney test, U = 3975, Z = -6.489, P < 0.0001.

### Dopamine D1 receptor activation enhances total protein synthesis in cultured cortical neurons

In light of our immunocytochemistry results, we further asked whether the dopamine D1 receptor-dependent eEF2 dephosphorylation coincides with enhanced protein synthesis. For this purpose, we utilized SUnSET, a nonradioactive method of monitoring global protein synthesis in cultured cells that uses puromycin to tag nascent proteins (Schmidt et al., 2009). Time-course experiments showed that puromycin incorporation was significantly increased after 1.5 and 4 hours of SKF38393 treatment (Figure 6A). In addition to measuring total levels of mRNA translation, we measured expression levels of BDNF and synapsin 2B proteins, which are known to be induced by D1 receptor and affected by eEF2 activity (Chong et al., 2006, Williams and Undieh, 2009). Prolonged stimulation of the cells with SKF38393 resulted in increased levels of BDNF and synapsin 2B after 4 hours of incubation (Figure 6B and Figure 6C).

**Figure 6.**
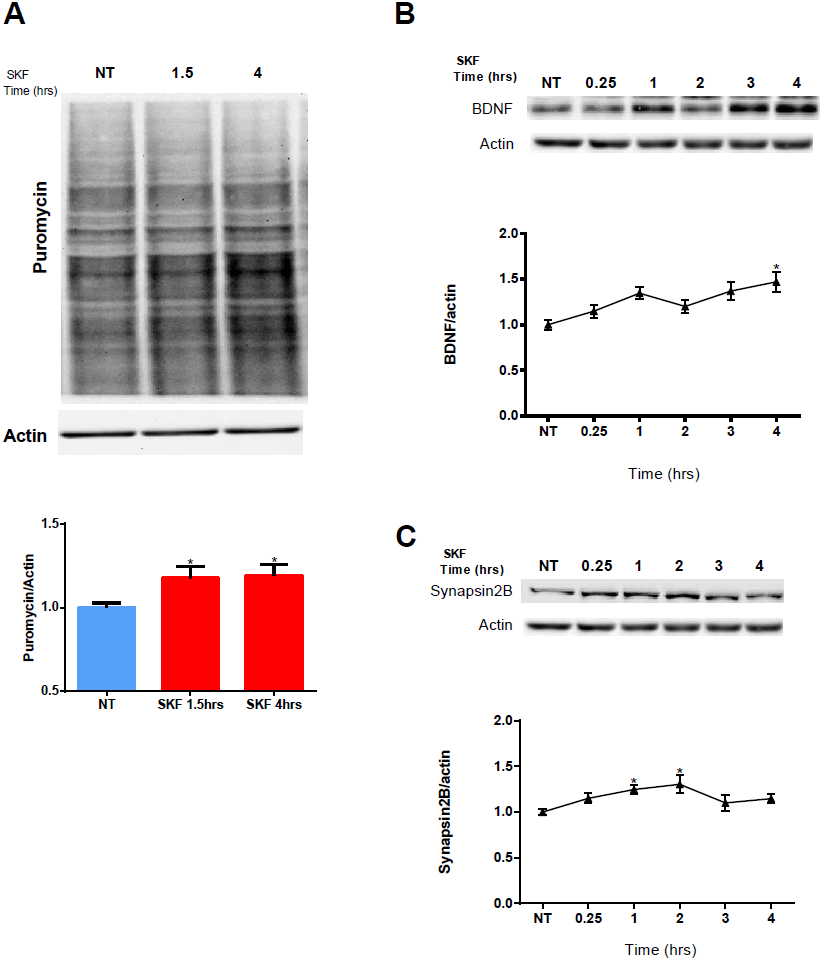
D1 receptor activation promotes general protein synthesis. **(A)** Representative Western blots and quantification of proteins labeled with puromycin from primary cortical neurons treated with SKF38393 (25μM) for 1.5 or 4 hours. Puromycin signal was normalized to p-actin loading control and quantified. **(B and C)** Levels of synapsin 2B and BDNF were quantified in primary cortical neurons treated with SKF38393 for the indicated time periods. Protein levels were normalized to β-actin loading control. Data are means ± SEM of five independent cultures. ^*^p<0.05. See also Figure 6-source data 1and Figure 6-figure supplement 1. **Source data 1-Statistical analysis of Figure 6** **Figure 6A**. One way ANOVA F_(2,37)_=5.354, p=0.009. Post-hoc test compared to Control; SKF1.5h, p=0.04; SKF4h, p=0.01. **Figure 6B.** One way ANOVA F_(5,29)_=5.756, p=0.001. Post-hoc test compared to Control; SKF1h, p=0.02; SKF3h, p=0.01; SKF4h, p=0.001. **Figure 6C.** One way ANOVA F_(5_,_29_)=3.744, p=0.01. Post-hoc test compared to Control; SKF1h, p=0.04; SKF2h, p=0.0.008.

The mTOR cascade promotes cap-dependent translation by phosphorylation and inhibition of 4E-BP (4E-binding protein) (E. Santini and E. Klann, 2011). Since D1 receptor stimulation induces phosphorylation of S6K (Figure 3B and Figure 4B), a well-known mTOR target (M.D. Antion, et al 2008; Hoffer, K.K et al 2011), we tested whether the increase in protein synthesis following SKF38393 treatment could also be a result of 4E-BP phosphorylation. Treatment of cortical neurons with SKF38393 did not change 4E-BP phosphorylation. However, rapamycin treatment did, as expected (Figure 6 supplement 1 A and Figure 6 supplement 1 B), suggesting that the D1 receptor-dependent increase in protein synthesis is mediated by eEF2 dephosphorylation.

In addition to its negative regulators (MEK-ERK and mTOR), eEF2K has also a positive regulator which is downstream to the D1 receptor, AMP kinase (AMPK). Examining the phosphorylation of AMPK at its activation site, Thr^172^ revealed no difference in the phosphorylation state at the same time points when D1 receptor inhibits eEF2K at Ser^366^ and leads to eEF2 dephosphorylation (15 and 60 min). This suggests that MEK-ERK and mTOR are indeed the dominant pathways for D1 receptor-dependent eEF2 dephosphorylation and increased protein synthesis.

### D1 receptor activation increases protein synthesis in neurons by eEF2K inactivation

To test directly if D1 receptor activation increases protein synthesis in an eEF2K/eEF2 pathway-dependent manner, we used primary cultures from eEF2K-KO mice. These mice show no eEF2 phosphorylation, but display normal phosphorylation of other translation factors (Adaikkan et al., 2018). Cortical cultures from eEF2K-KO mice and their wild-type littermates were treated for 4 hours with SKF38393 and global protein synthesis was analyzed by SUnSET. Immunocytochemistry and western blot analyses of puromycin incorporation showed a significant increase in wild type but not in eEF2K-KO cultures following SKF38393 incubation (Figure. 7A and Figure 7B, Figure 7 supplement 1 A-Figure 7 supplement 1 C). To probe whether the D1 receptor activation-induced increase in translation is dependent on ERK/mTOR/eEF2K/eEF2 signaling, we pre-treated cortical neurons derived from wild type and eEF2-KO mice with or without MEK inhibitor U0126, followed by SKF38393 treatment. We found that the SKF38393-mediated increase in translation was reduced by treatment with U0126 in wild type neurons, while no change in puromycin incorporation was detected in eEF2K-KO mouse-derived cultures treated with U0126. These results suggest that the MEK/ERK and eEF2 pathways mediate the D1 receptor-dependent increase in protein synthesis in cortical neurons.

**Figure 7.**
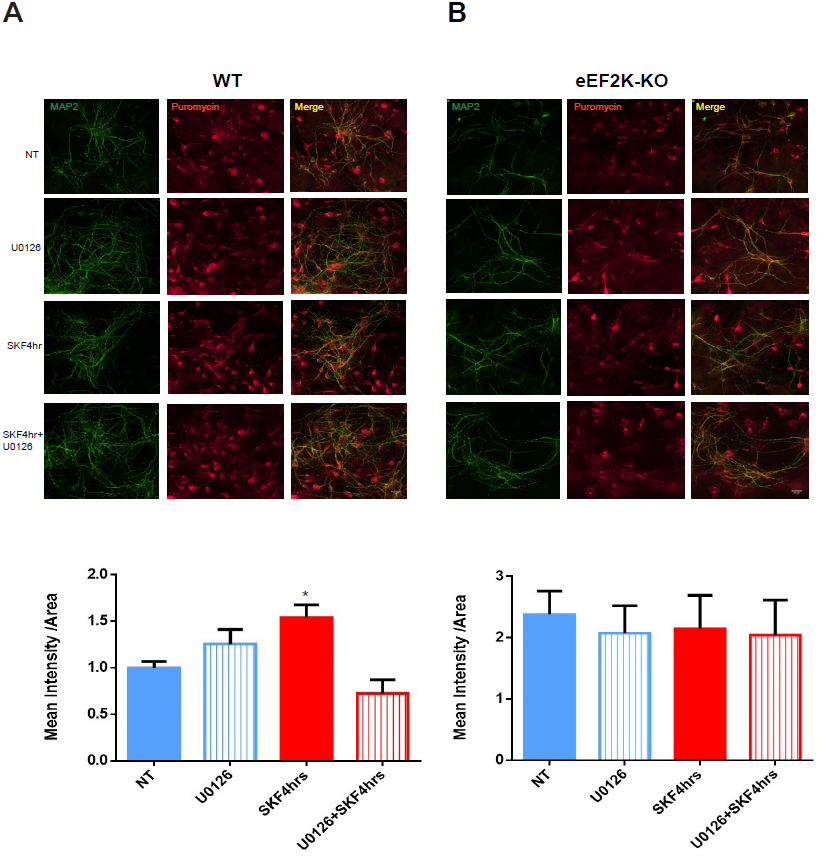
Inhibition of MEK/ERK pathway attenuates D1 receptor-dependent increase in protein synthesis in WT but not eEF2-KO primary cultures. **(A and B)** Cortical primary cultures from WT or eEF2K-KO mice were incubated with SKF38393 (25μM) for 4 hours with or without pre-incubation with MEK inhibitor U0126 (20μM) for 30 min, followed by incubation with puromycin (1μg/ml) for 10 min. Puromycin was detected by immunofluorescence, and quantification of puromycin labeling (red) was done by measuring mean intensity of MAP2 (green) positive cells. Data are means ± SEM of four independent cultures for wild-type and three independent cultures (^*^p<0.05). See also Figure 7-source data 1 and Figure 7-figure supplement 1. **Source data 1-Statistical analysis of Figure 7** **Figure 7A.** One way ANOVA F _(3,17)_=7.909, p=0.002. Post-hoc test compared to Control; SKF4h, p=0.01; post-hoc for SKF4h compared to SKF4h+U0126, p=0.001. **Figure 7B.** One way ANOVA F _(3,8)_=0.176, p=0.91.

## Discussion

In summary, our data show that dopamine D1 receptor activation modulates eEF2 phosphorylation and increases protein synthesis in primary cortical neurons. We found that NMDA receptor is required for the D1 receptor-dependent induction of MEK/mTOR activity that leads to inactivation of eEF2K. The reduction in eEF2K activity is sufficient to induce a marked eEF2 dephosphorylation in neurons, especially in dendrites, and increased global mRNA translation.

These data support the view that dopamine D1 receptor activation influences protein synthesis, an important component of memory and synaptic plasticity formation (Andre and Manahan-Vaughan, 2015, Fenu et al., 2001, Hikind and Maroun, 2008, Nader and LeDoux, 1999, Roffman et al., 2016, Schicknick et al., 2012). In addition, these data provide insight into the kinetics and mechanism of the signaling pathways involved in regulation of eEF2K activity in neurons. Our data provide further decoding of the well-known relationship between D1 and NMDA receptors and its effect on signal transduction (David et al., 2014, Dunah and Standaert, 2001, Lee et al., 2002, Martina and Bergeron, 2008, Pei et al., 2004, Stramiello and Wagner, 2008). Finally, our results indicate that, in addition to the canonical calcium-calmodulin-dependent activation of eEF2K following NMDA receptor stimulation, NMDAR-D1 partnership induces translational changes via ERK and S6K signaling cascades. This includes eEF2 dephosphorylation in neurons, in the soma and especially in dendrites, which is likely to contribute to the observed increase in polypeptide elongation and expression of BDNF and synapsin 2B (Adaikkan et al., 2018, Autry et al., 2011, Belelovsky et al., 2009, Gildish et al., 2012, Im et al., 2009, Ma et al., 2014, Park et al., 2008, Taha et al., 2013). The data obtained using cortical cultures derived from eEF2K-KO mice led to the notion that the eEF2K pathway accounts for the increase in protein synthesis following dopamine D1 receptor activation.

eEF2K has been identified as a biochemical sensor that is tuned to the pattern of neuronal stimulation (Heise et al., 2014, McCamphill et al., 2015, Sutton et al., 2007). Perturbation of intracellular calcium concentration by NMDA and neuronal activity produces eEF2K/eEF2-dependent changes of dendritic proteome (Ehlers, 2003, Lazarevic et al., 2011, Perez-Otano and Ehlers, 2005, Turrigiano, 2008, Turrigiano and Nelson, 2004, Virmani et al., 2006). Although eEF2K activity is controlled principally by Ca^+2^-Calmodulin complex, it is also regulated by phosphorylation. mTOR- and MEK-dependent phosphorylation of eEF2K reduces its activity, while PKA and AMPK mediated phosphorylation does the opposite (Heise et al., 2014, Taha et al., 2013, Wang et al., 2001). Given this complex regulation of its function, it is tempting to speculate that eEF2K works as a signaling hub, linking synaptic activity to protein synthesis. Conversely, eEF2 dephosphorylation occurs by spaced 5-HT activity and the PKA stimulation (McCamphill et al., 2015). eEF2K activity can be regulated by the type of neurotransmission in which eEF2K acts as a biochemical sensor to discriminate between evoked action potential and spontaneous miniature synaptic transmission (Sutton 2007).

In a similar manner, optimal levels of dopamine D1 receptor stimulation in dendrites work to gate out “noise”, while high levels, e.g., during stress, suppress delay firing (Arnsten et al., 2015). For instance, maintenance of synaptic strength in hippocampal slices treated with low concentrations of dopamine D1 agonist requires MEK and CaMKII activation, while in slices treated with high concentrations maintenance of synaptic strength is dependent only on MEK activation (Barcomb et al., 2016). Therefore, it is not surprising that dopamine D1 receptor agonist serves potentially as an antidepressant (D’Aquila et al., 1994). This effect may be due to the ability of dopamine to regulate neuronal function via the eEF2K pathway.

On the other hand, mRNA translation following synaptic activation can also be regulated by other pathways such as the mTOR/4E-BP pathway (Scheetz et al., 2000). It was shown that, in primary striatal neurons, the antipsychotic drug haloperidol activates translation-related pathways mediated by mTOR, which results in increased phosphorylation of 4E-BP and S6K1 (Bowling et al., 2014). Moreover, our results show that the phosphorylation status of 4E-BP does not change following D1 receptor agonist treatment, but it does affect the eEF2K pathway, which leads to D1 receptor-dependent increased protein synthesis. Recent studies involving manipulation of eEF2K activity have been a field of intense research, ranging from addiction (Sutton and Caron, 2015, Werner et al., 2018) and depression. Indeed, eEF2K serves as a target for the antidepressant ketamine, and has a role also in the progression of neurodegenerative diseases (Adaikkan et al., 2018, Heise et al., 2017, Jan et al., 2018, Kokkinou et al., 2018, Zang et al., 2018).

Our findings establish a connection between dopamine D1 receptor activation and eEF2K, and open a door to better understanding the molecular mechanism underlying the role of neurotransmitters in synaptic plasticity, addiction, and antidepressants. Future studies using neurodegenerative disease animal models, combined with pharmacological and behavioral paradigms will better link the eEF2 pathway and a potential target for therapy.

## Material and Methods

### Mice

eEF2K-KO mice, in which coding exons 7, 8, 9, and 10 of eEF2K were deleted, were generated by the laboratory of Christopher G. Proud. We derived eEF2K wild-type (WT) and KO littermates by crossing heterozygous mice as previously described (Heise et al., 2017), eEF2K mice were bred from colonies maintained at the University of Haifa. C57BL/6 mice were obtained from local vendors (Envigo RMS, Jerusalem, Israel) and after acclimation to the facility were used for experiments. Animals were provided ad libitum with standard food and water and were maintained on a 12/12 h light/dark cycle. All experiments were approved by the Institutional Animal Care and Use Committee of the University of Haifa, and adequate measures were taken in order to minimize pain, in accordance with the guidelines laid down by the European Union and United States NIH regarding the care and use of animals in experiments.

### Cortical cell culture

Primary cortical neuronal cultures were isolated from P0 or P1 C57BL/6J or eEF2K-WT or KO mice of either sex as previously described (Ounallah-Saad et al., 2014). Briefly, both hippocampi were removed, and cortical regions were taken. The tissue was chemically dissociated by trypsin and DNase, and mechanically, using a siliconized Pasteur pipette. Cells were plated onto round coverslips coated with 20μg/ml Poly-L-ornithine and 3 μg/ml laminin (Sigma), placed in 6-well plates (300,000 cells per well) or 12-well plates (150,000 cells per well). Culture medium consisted of MEM (Gibco), 25 μg/ml insulin (Sigma), 27.8 mM glucose (Sigma), 2 mM l-glutamine (Sigma), and 10% horse serum (Biological Industries, Israel). Cultures were maintained at 37°C in a 95% air/5% CO_2_ humidified incubator. Half the volume of the culture medium was replaced at days 8 and 11 with feeding medium containing glutamine 2mM, insulin 25 μg/ml, and 2% B-27 supplement (Gibco).

### In vivo experiments: Injection of mice with SKF38393

Adult male C57BL/6 mice were injected i.p. with either 0.5mg/Kg SKF38393 or vehicle (0.9% NaCl saline) and returned to their cage. The animals were sacrificed 15 minutes later by cervical dislocation, the prefrontal cortex and the hippocampus were collected on ice, and the tissue was immediately frozen in liquid nitrogen. The tissue was stored at -80°C until further use.

### Collection of tissue samples

The obtained tissue samples were homogenized using a glass-Teflon homogenizer in lysis buffer containing 10 mM HEPES, pH 7.4, 2 mM EDTA, 2mM EGTA, 0.5mM DTT, 1% phosphate inhibitor cocktail (Sigma), and 1% protease inhibitor cocktail (Sigma). Protein quantification was done using a BCA protein assay kit (GE Healthcare). Appropriate volumes of 2X SDS sample buffer (10% glycerol, 5% β-mercaptoethanol, 4% SDS, 120 mM Tris-HCl, pH 6.8) were added to the homogenates, and samples were boiled for 5 min and stored at -20°C.

### Pharmacological treatments

After 14 days in vitro cortical neurons were treated with dopamine D1 receptor agonist SKF38393 (25μM, Sigma) or the D2 agonist quinpirole (10μM, Sigma) in a time-dependent manner. Cells were pre-incubated with the following drugs for 30 minutes before agonist treatment as indicated: D1 receptor antagonist: SCH23390 (10μM, Sigma); D2 receptor antagonist: eticlopride (20μM, Sigma); mTORC1 inhibitor: rapamycin (100nM, Sigma); MEK inhibitor: U0126 (20μM); NMDAR antagonist: APV (40μM). After incubation with the antagonists, cells were treated with SKF38393 (25μM) for the indicated time periods.

### Immunocytochemistry

Primary cortical cultures were fixed in cold 4% formaldehyde solution in phosphate saline buffer 0.01M (PBS) for 10 min. The cells were washed three times for 5 min with PBS + Triton X-100 1%. The cells were then incubated for 1 hour at room temperature (RT) in blocking solution of PBS + Triton ×-100 1%, containing 10% fetal calf serum and 0.3% bovine serum albumin (BSA).

The cells were incubated overnight at 4°C and 1 hour at room temperature (RT) with the following primary antibodies diluted in the same blocking solution: phospho-eEF2 Thr^56^ (1:100, Cell Signaling), eEF2 (1:100, Cell Signaling), MAP2 (1:1000 Abcam), puromycin (1:1000, Millipore). Cells incubated in blocking solution lacking the primary antibody were used as negative control. After washing with PBS + 1% Triton (3 × 5min), cells were incubated for 1 hour at RT with the corresponding secondary antibodies: donkey Anti-Chicken AlexaFluor^®^ 488 (1:500), donkey Anti-Rabbit and AntiMouse AlexaFluor^®^ 594 (1:500). The cells were then washed with PBS + 1% Triton (2 × 5 min) and PBS (2 × 5 min). Finally, coverslips were mounted on Superfrost^TM^ Plus Adhesion slides (Thermo Scientific) with Slow Fade^®^ Gold antifade reagent containing DAPI (Life Technologies).

### Immunocytochemistry quantification

Signal intensity quantification of phosphorylated eEF2 (peEF2) in the cell soma or dendrites was done by NIS Element Advanced Research (Ar) 4.5 (Nikon Japan) software in MAP2 labeled neurons. Confocal images were acquired using a Nikon 63X immersion oil objective at a resolution of 1024×1024 pixels. Each image was a Z series projection of 7 to 10 images, taken at depth intervals of 0.5 μm. To define the region of interest for quantification, cell bodies and dendrites close to the cell body were manually traced using NIS Element AR software on the MAP2 channel. peEF2 signal intensity (mean pixel intensity) was estimated as the peEF2 integrated fluorescence intensity divided by the area marked by the MAP2 signal.

### Puromycin immunocytochemistry and quantification

Cells for immunofluorescence were plated on coverslips coated with Poly-L-ornithine and laminin coating. After 14 DIV cells were pre-treated with U0126 (20 μM) or vehicle (DMSO) for 30 minutes and then treated with SKF38393 (25 μM) for 4 hours. After SKF38393 treatment, cells were further incubated with 10μg/ml puromycin for 10 min in the same medium and fixed in 4% paraformaldehyde. Cells were stained with anti-puromycin (1:1000) and MAP2 (1:1000) antibodies, following the same procedure for peEF2Thr^56^ immunocytochemistry. Images were taken as Z series projection of 5 to 9 images at depth intervals of 0.25 μm at ×60 magnification with an Olympus IX81 microscope using Olympus cellSens1.16 software. Quantification of puromycin incorporation was done by selecting neurons randomly in MAP2 labeled neurons and estimating the puromycin signal as mean intensity divided by the area marked by MAP2 signal using ImageJ 1.51J software. Quantification was done in a blind manner based on three independent experiments for each condition.

### SUnSET

Protein synthesis was measured by the SUnSET method. Cortical neurons were isolated and maintained for 14 days in culture, as described above. Neurons were pre-treated with U0126 (20 μM) or vehicle (DMSO) for 30 minutes and then treated with SKF38393 (25 μM) for 1.5 or 4 hours. After SKF38393 treatment, cells were further incubated with 10μg/ml puromycin for 10 min in the same medium. After puromycin labeling, cells were washed once with PBS and lysed in homogenization buffer (40mM Tris-HCl, pH 8.0, 150mM NaCl, 25mM β-glycerophosphate, 50mM NaF, 2mM Na_3_VO_3_, 10% glycerol, 1% Triton X-100). Puromycin incorporation was detected by Western blotting using 12 D 10 antibodies for puromycin (1:5000, Millipore).

### Western blotting

Samples in SDS sample buffer were subjected to SDS-PAGE (7.5-10%) and Western blot analysis. Lanes were loaded with an equal amount of protein. Following transferring into a nitrocellulose or PVDF membranes using TransBlot^®^ Turbo^TM^ Transfer System (Bio-Rad), bands were visualized with Ponceau staining (Bio-Rad). Membranes were blocked in 5% BSA or 5% nonfat-dry milk (depending on the primary antibody) for 1 hour at RT, before being incubated overnight at 4°C with the primary antibodies: p44/42 MAP Kinase (1:1000, Cell Signaling) and Phospho-P44/42 MAP Kinase-(Thr^202^/Tyr^204^) (1:1000, Cell Signaling); S6K (1:1000, Cell Signaling), phospho-S6K(Thr^389^) (1:750, Cell Signaling), eEF2 (1:1000, Cell Signaling), phospho-eEF2(Thr^56^) (1:1000, Cell Signaling), phpho-eEF2K(Ser^366^), BDNF (1:500 Santa Cruz), synapsin 2B (1:1000 Abcam), β-Actin (1:6000, Abcam). AMPK (Cell Signaling), phosphor-AMPK (Thr172), 4E-BP(1:1,1000, Cell Signaling), phosphor-4E-BP (Thr^37^/^46^) (1:1,000, Cell Signaling).

Following three 5-min washing steps in Tris-buffered saline (140 mM NaCl, 20 mM Tris, pH 7.6) plus 0.1% Tween 20 (TBST), membranes were incubated for 1 h at room temperature with secondary HRP-linked antibodies: Goat-anti-Rabbit (IgG) HRP conjugated; Goat-anti-chicken (IgG) HRP conjugated (1:10,000) (Jackson ImmunoResearch). Immunodetection was accomplished with the Enhanced Chemiluminescence EZ ECL kit (Biological Industries). Quantification of immunoblots was performed with a CCD camera and Quantity One 4.6 software (Bio-Rad). Each sample was measured relative to the background. Phosphorylation levels were calculated as the ratio of phosphorylated protein and a total amount of protein.

### Statistical analysis

Graphs were prepared using GraphPad Prism 6.01, InStat Software (GraphPad Software, CA, USA). Data are expressed as mean ± SEM. Statistical analysis was performed using SPSS version 24. Statistical significance was determined with one-way ANOVA followed by Tukey’s post hoc test. Student’s t-test or Mann-Whitney were used to examine the differences between groups.

**Figure 8.**
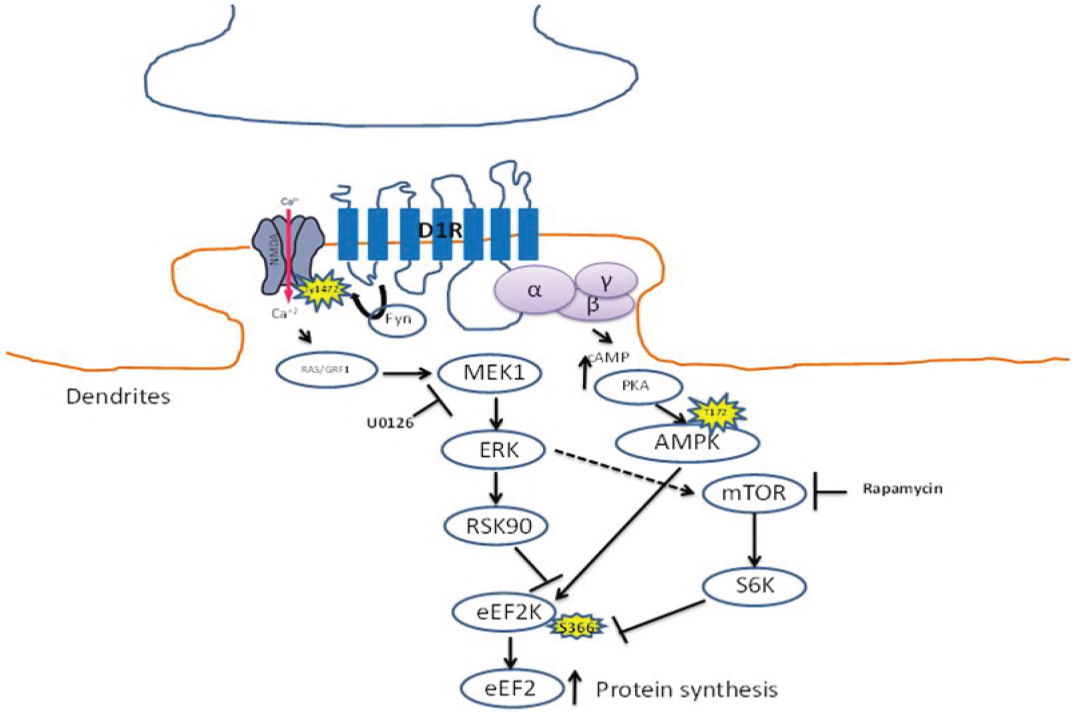
Model of D1 receptor-dependent dephosphorylation of eEF2 in cortical neurons. Dopamine D1 receptors activated in dendrites lead to small calcium influx via the NMDA receptor, which elevates both MEK–ERK and mTOR pathways. Both pathways inhibit eEF2K activity by phosphorylating its Ser^366^ site, leading to eEF2 Thr^56^ dephosphorylation and increased local protein synthesis.

## Acknowledgments

We thank Prof. Christopher G. Proud (Australian Health and Medical Research Institute, Adelaide, Australia) for providing us with the eEF2K-KO mice. This research was supported by a grant from the Canadian Institutes of Health Research (CIHR), the International Development Research Centre (IDRC), the Israel Science Foundation (ISF) and the Azrieli Foundation (ISF- IDRC 2395/2015); ISF 946/17; ISF-UGC 2311/15; and TransNeuro ERANET JPND supported by the Israel Ministry of Health Grant 3-14616 to K.R.

---

**Figure 1 supplement 1.**
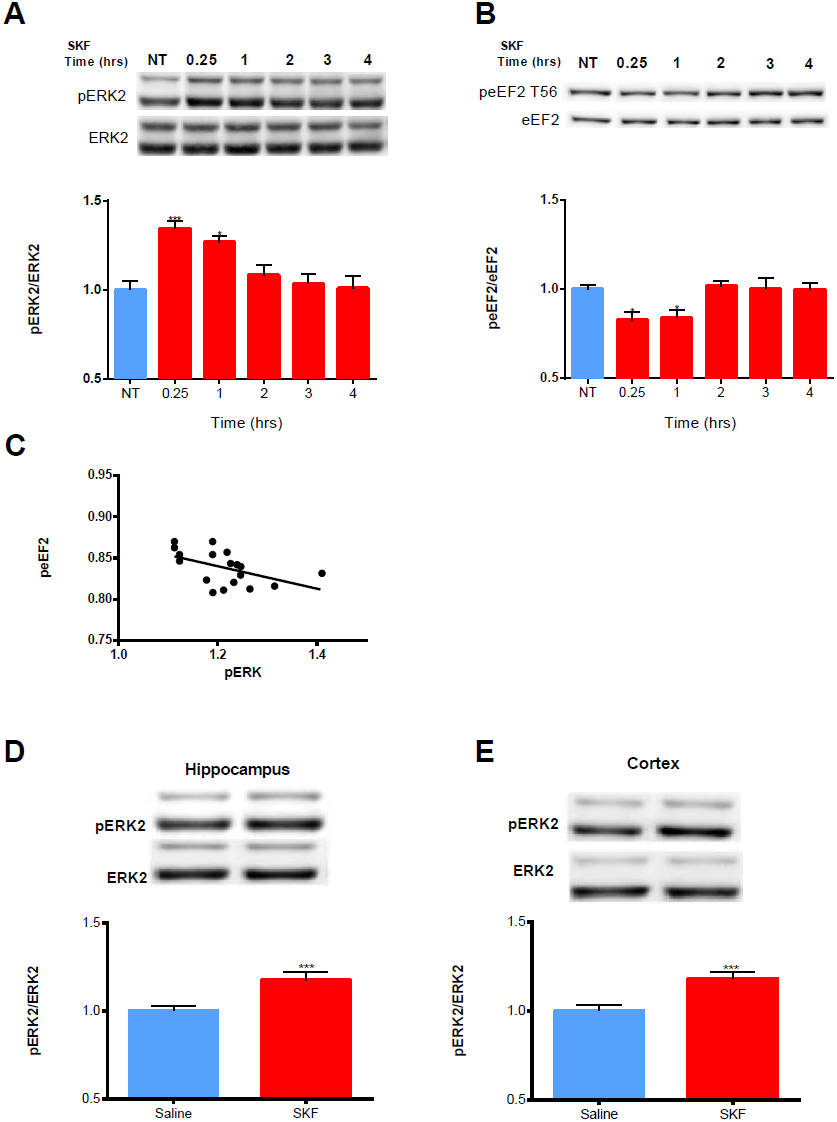
D1 receptor activation induces ERK1/2 phosphorylation both in vitro and in vivo and correlates with eEF2 dephosphorylation. (A)Representative blots and quantification of the ratios of phosphorylated to total ERK1/2 in cortical primary cultures incubated with D1 receptor agonist SKF38393 (25μM) for the indicated time points (from five independent cultures). One way ANOVA F_(5,29)_=11.912, p<0.0001. Post-hoc test compared with control: SKF 15min, p<0.0001; SKF60min, p<0.0001. **(B)** Western blot analysis of eEF2 (Thr^56^) phosphorylation in cortical primary cultures treated as in A. One way ANOVA F(_5,29)_=6.338, p<0.0001. Post-hoc test compared with control: SKF15min, p=0.006; SKF60min, p=0.01; post-hoc for SKF15min compared with: SKF2h, p=0.001; post-hoc compared to 3h, p=0.02; compared to 4h, p=0.02; post-hoc for SKF60min: compared to SKF2h, p=0.01; compared to SKF3h, p=0.03; compared to SKF4h, p=0.04. **(C)** Pearson’s correlation of eEF2 phosphorylation levels with ERK2 phosphorylation levels. Pearson’s correlation: r=-0.50, p<0.05. **(D and E)** Representative Western blots and quantification of the ratio of phosphorylated to total ERK1/2 in hippocampal and cortical extracts from C57BL/6 male mice injected i.p. with SKF38393 (5mg/Kg) and sacrificed 15 minutes later. Data are means ± SEM of five independent cultures ^***^p<0.001. Hippocampus: Independent samples t-test: saline, n = 12 mice; SKF38393, n=12 mice; t (_22_) = - 3.087, p=0.005; cortex: Independent samples t-test: saline, n = 12 mice; SKF38393, n=12 mice; t _(29)_ =-4.199, p<0.0001. SKF, SKF38393.

**Figure 1 supplement 2.**
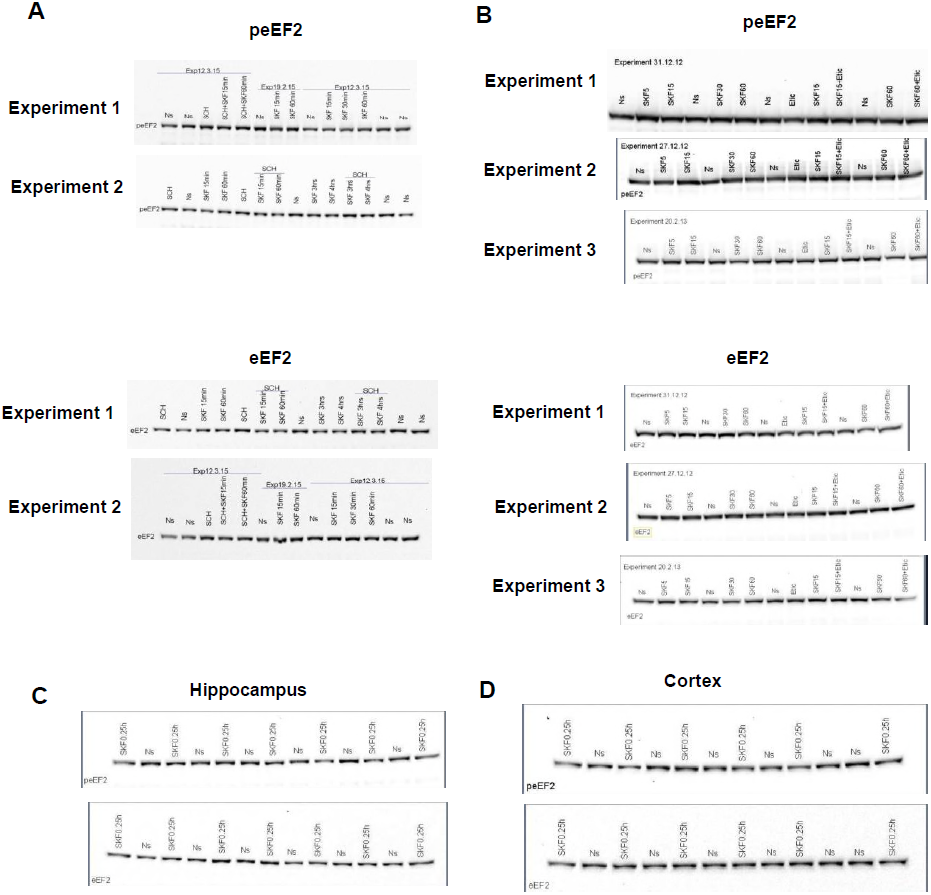
Uncropped original immunoblots of Figure 1. **(A)** Original immunoblots of peEF2 and eEF2 for Figure1A following incubation with SKF38393 with/without SCH23390 from 3 independent experiments. **(B)** Original blots of peEF2 and eEF2 for Figure 1B following incubation with quinpirole with/without eticlopride from 3 independent experiments. **(C,D)** Original immunoblots for peEF2 and eEF2 from hippocampus and cortex of 6 C57BL/6 mice in Figure 1C,D.

**Figure 1 supplement 3.**
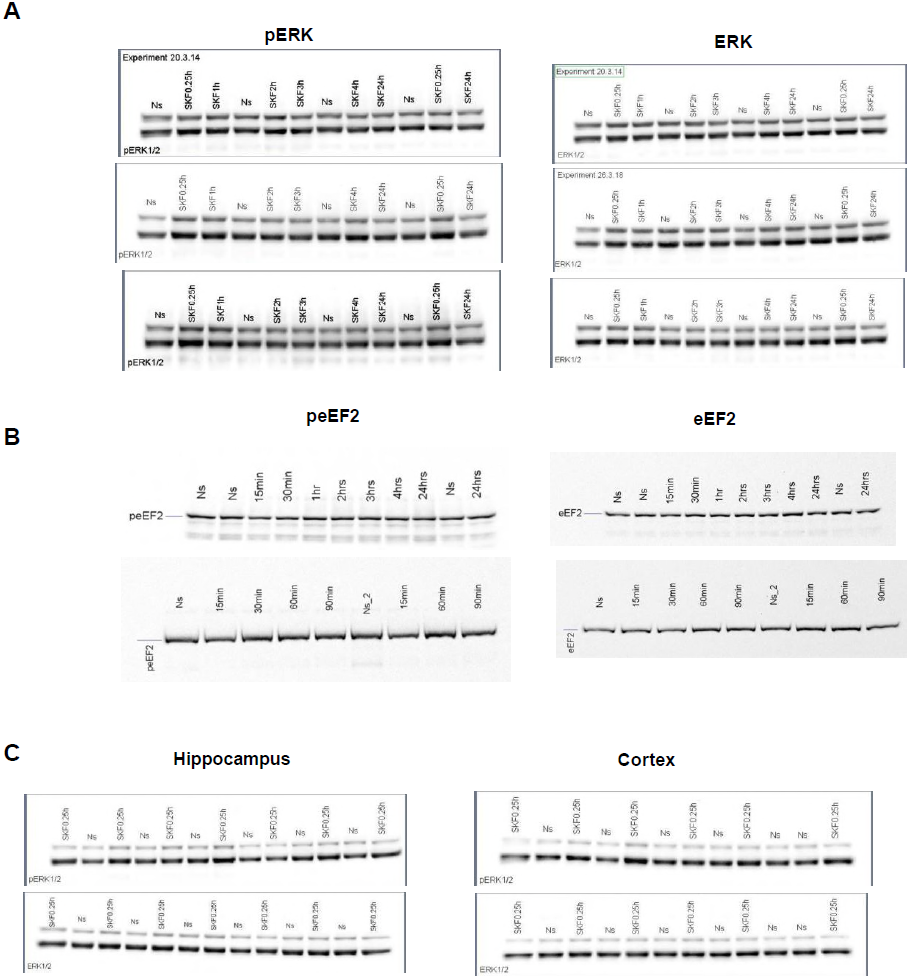
Uncropped original immunoblots of Figure supplement 1. **(A)** Original immunoblots of pERK1/2 and ERK1/2 for Figure supplement 1A following prolonged incubation with SKF38393 (up to 24h) from 3 independent experiments are shown. **(B)** Original blots of peEF2 and eEF2 for Figure 1 supplement B following prolonged incubation with SKF38393 (up to 24h) from 3 independent experiments. **(C,D)** Original immunoblots for pERK1/2 and ERK1/2 from hippocampus and cortex of 6 C57BL/6 mice in Figure supplement 1C,D. The same samples as in Figure 1 C,D were used in these blots.

**Figure 2 supplement 1.**
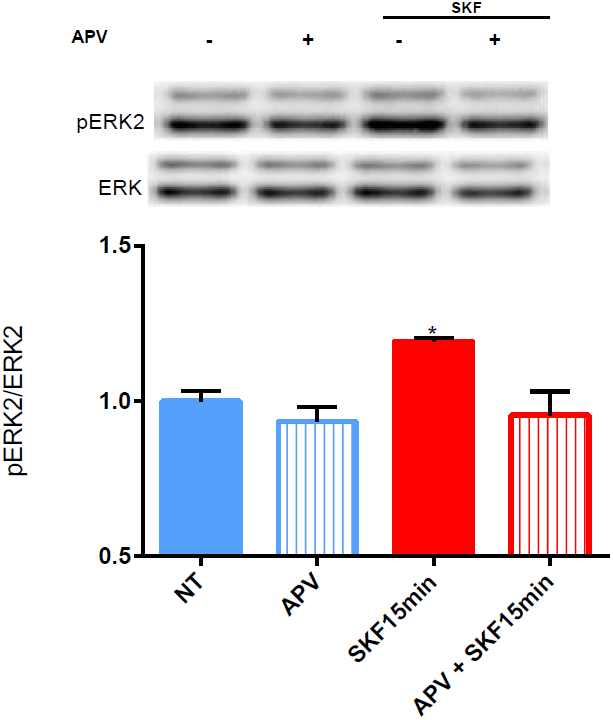
ERK1/2 phosphorylation by the D1 stimulation requires the NMDA receptor. (A) Representative blots and quantification of the ratios of phosphorylated ERK2 (Thr^202^/Tyr^204^) to total ERK1/2 in cortical primary cultures incubated with SKF38393 (25μM) for 15 min with or without pretreatment of the NMDA receptor antagonist APV (40μM). Data are means ± SEM of five independent cultures. One way ANOVA F_(3,26)_= 5.830, p=0.003. Post-hoc test compared to Control, SKF 15min, p=0.03; post-hoc for SKF 15min, compared to SKF 15min+APV, p=0.003.

**Figure 2 supplement 2.**
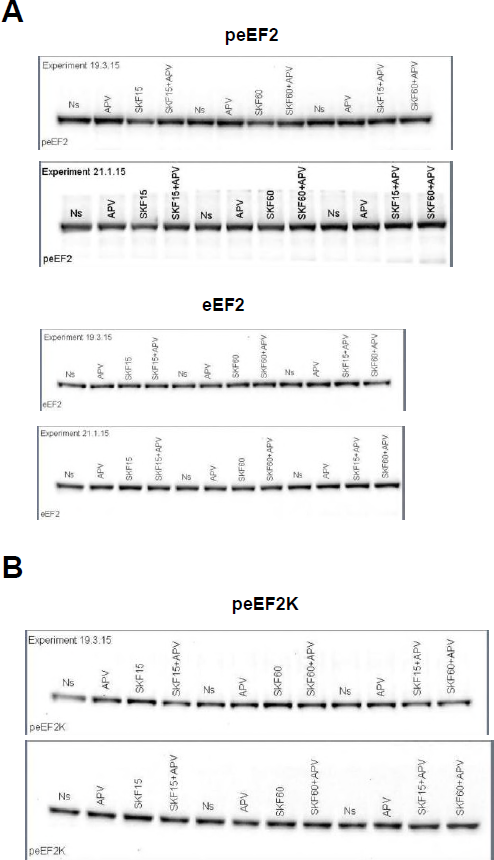
Uncropped original immunoblots of Figure 2. **(A)** Original immunoblots of peEF2 and eEF2 for Figure 2A following incubation with SKF38393 with/without APV from 3 independent experiments. **(B)** Original blots of peEF2K for Figure 2B.

**Figure 2 supplement 3.**
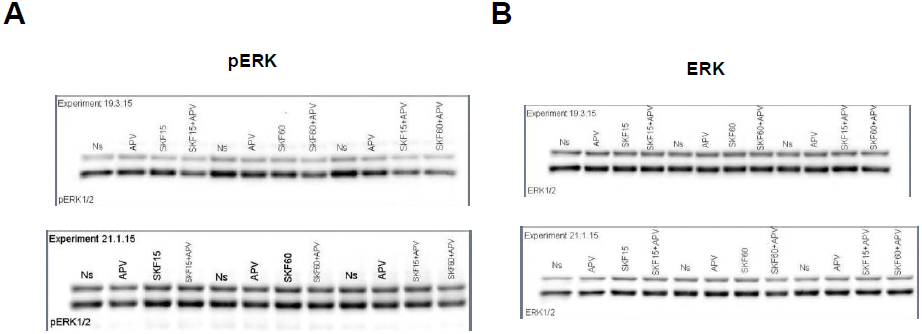
Uncropped original immunoblots of Figure supplement 2. **(A)** Original immunoblots of pERK1/2 and ERK1/2 for Figure supplement 2 following incubation with SKF38393 with/without APV. The blots correspond to the experiments shown in Figure 1.

**Figure 3 supplement 1.**
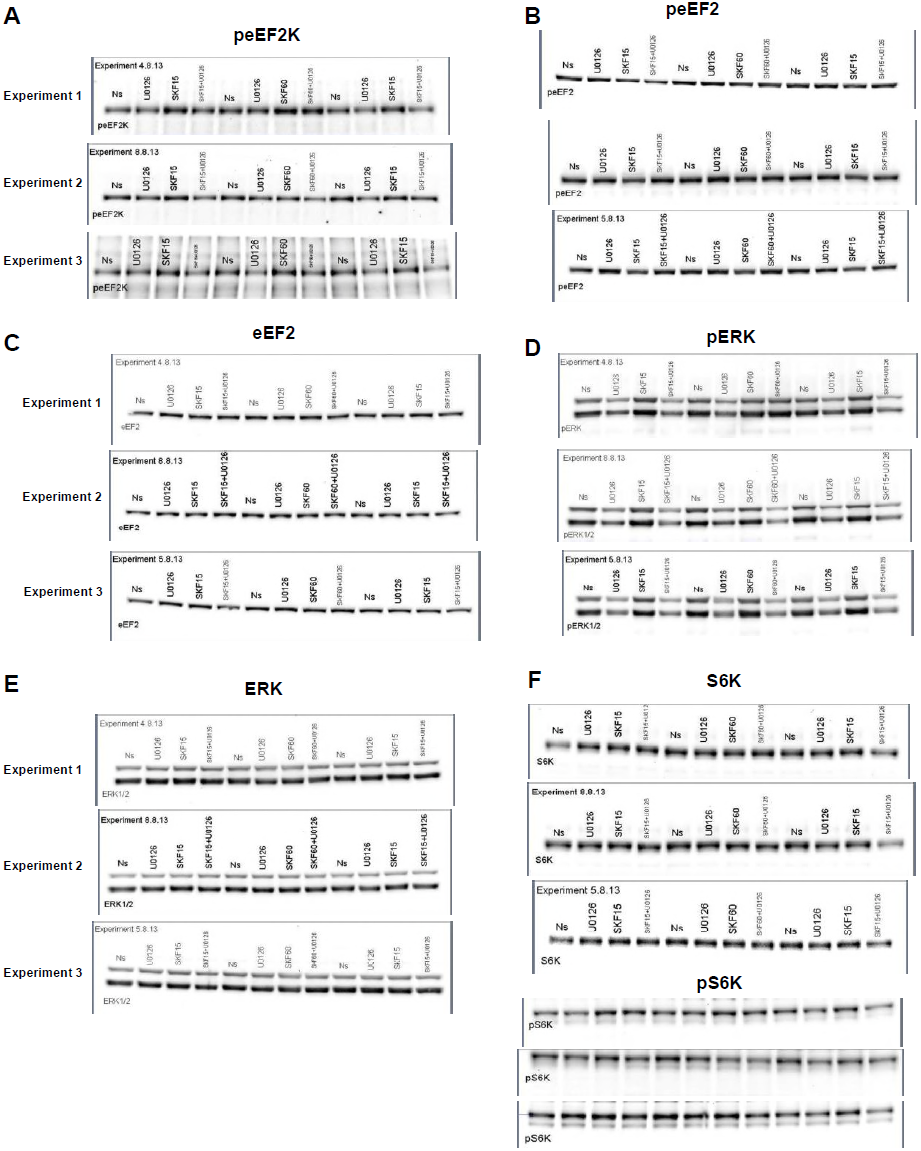
Uncropped original immunoblots of Figure 3. **(A)** Original immunoblots of peEF2K, peEF2, and eEF2 for Figure3A following incubation with SKF38393 (25μM), with/without pretreatment with MEK inhibitor U0126 (20μM) for 30 min 3 independent experiments are shown. **(B)** Original blots of pERK1/2, ERK1/2, pS6K, and S6K following SKF38393 (25μM), with/without pretreatment with MEK inhibitor U0126 (20μM) for 30 min of the experiments shown in Figure 3B.

**Figure 4 supplement 1.**
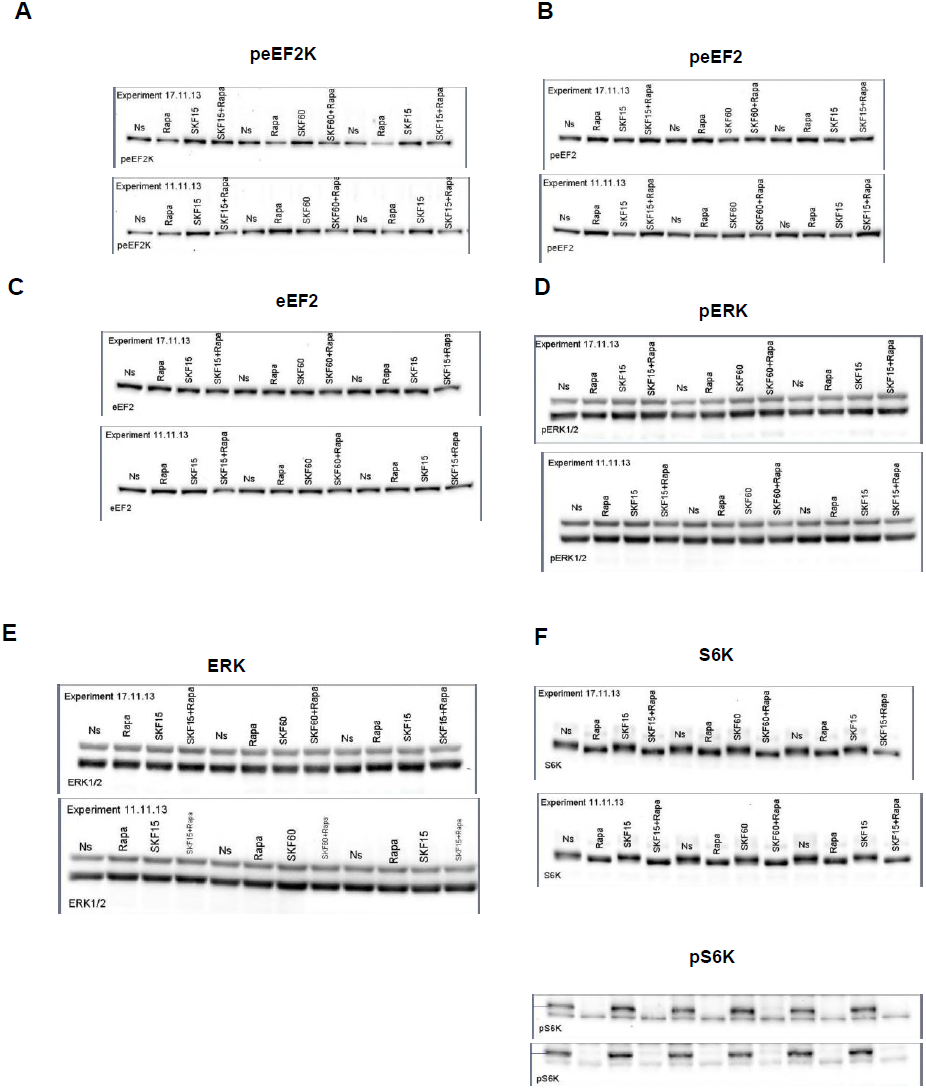
Uncropped original immunoblots of Figure 4. **(A)** Original immunoblots of peEF2K, peEF2, and eEF2 for Figure4A following SKF38393 (25μM) for 15 and 60 min with/without pretreatment of rapamycin (100nM) for 30 min from 3 independent experiments. **(B)** Original blots of pERK1/2, ERK1/2, pS6K, and S6K from the same experiments following SKF38393 with/without rapamycin for Figure 4B.

**Figure 5 supplement 1.**
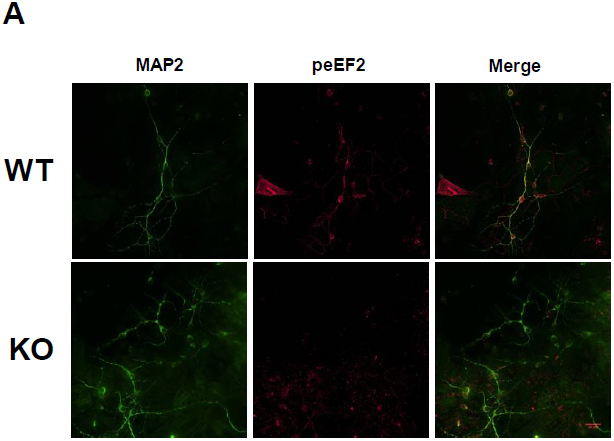
Representative images of phospho-eEF2 in primary cultures from WT and eEF2-KO mice. Immunofluorescence using phospho-eEF2 (red) and MAP2 (green) antibodies in cortical primary cultures from WT and eEF2K- KO mice. Scale bar 20μm.

**Figure 6 supplement 1.**
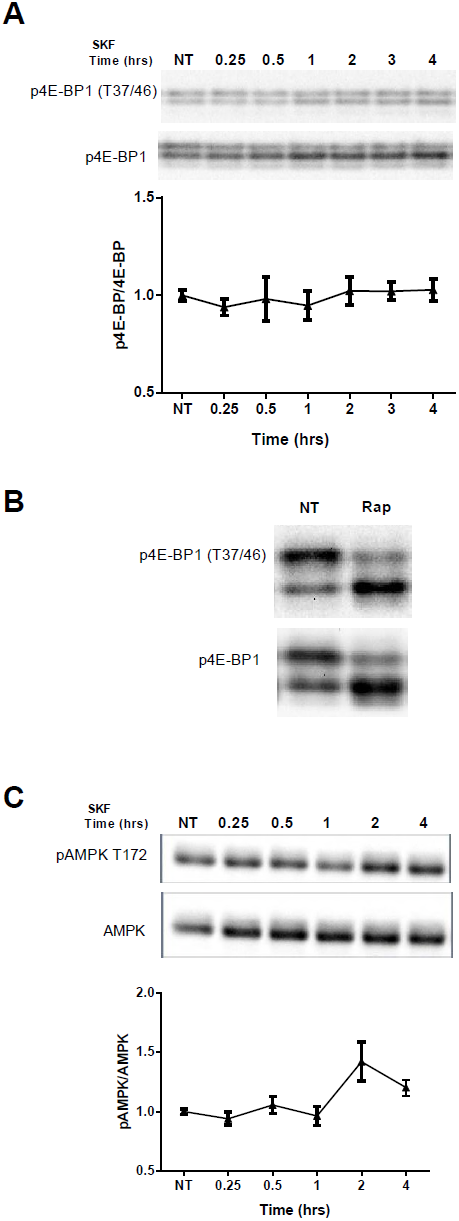
D1 receptor activation does not have an effect on 4E- BP phosphorylation and AMPK. **(A)** Representative blots and quantification of the ratios of phosphorylated 4E-BP (Thr^37/46^) to total 4E-BP from cortical primary cultures incubated with SKF38393 (25μM) and harvested for the indicated time periods. Data are means ± SEM of five independent cultures Data are means ± SEM of five independent cultures. One way ANOVA F_(6,30)_=0.365, p=0.895 **(B)** Representative control blots for phosphorylated 4E-BP1 (Thr^37/46^) and total 4E-BP1 from cortical primary cultures incubated with or without rapamycin (100nM). **(C)** Representative control blots for phosphorylated AMPK (Thr^172^) and total AMPK from cortical primary cultures incubated with SKF in the indicated time points. One way ANOVA F_(5,53)_=3.008, p=0.01.

**Figure 6 supplement 2.**
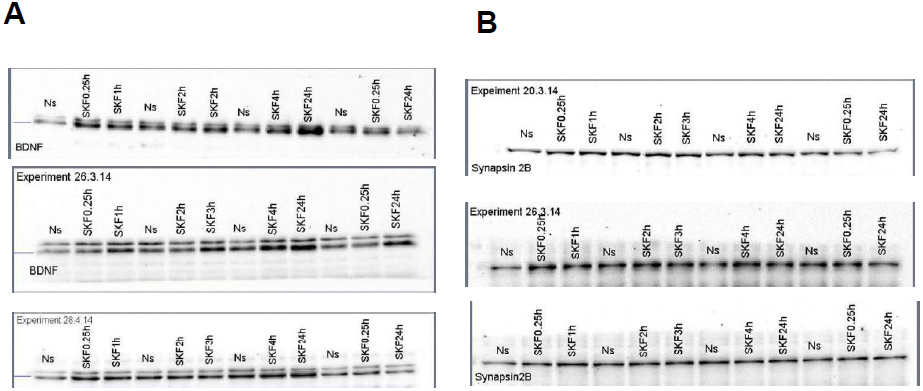
Uncropped original immunoblots of Figure 6. **(A,B)** Original immunoblots of BDNF and synapsin 2B following prolonged stimulation of SKF38393 (up to 24h) from three independent experiments for Figure 6A,B. The blots are from the same three experiments as in Figure supplement 1A (pERK, peEF2).

**Figure 6 supplement 3.**
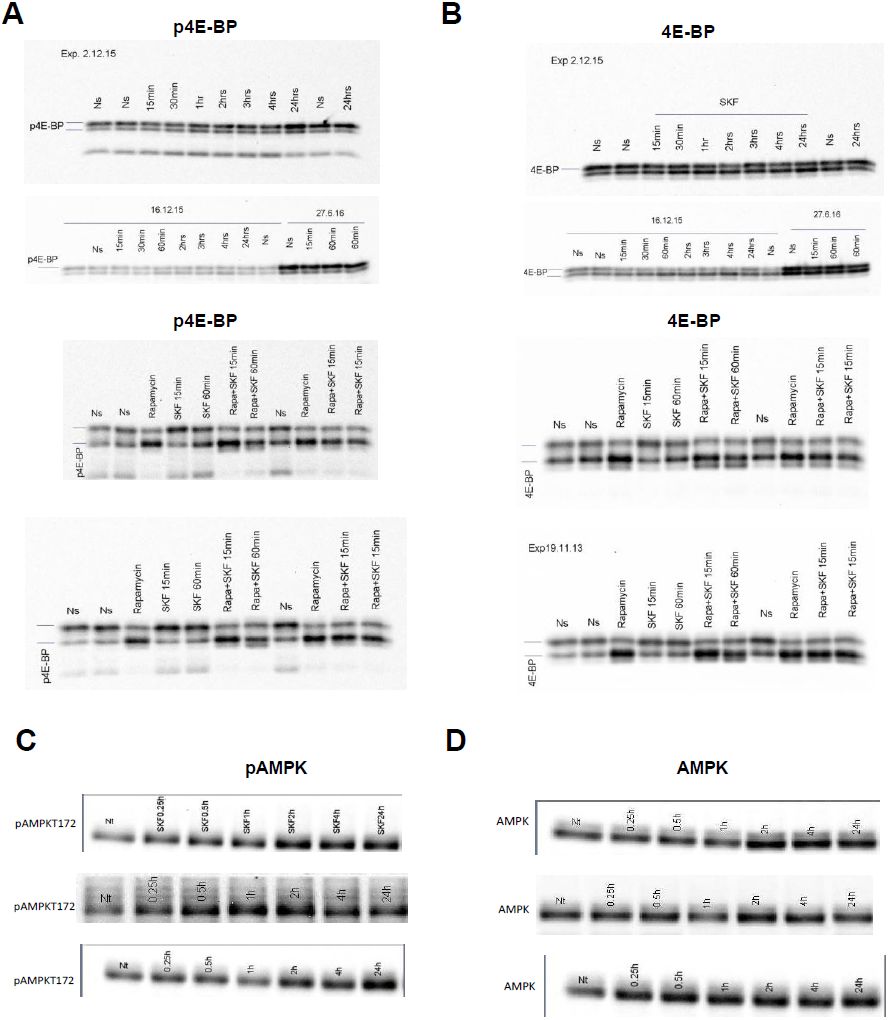
Uncropped original immunoblots of Figure supplement 1. **(A)** Original immunoblots of p4E-BP and 4E-BP for Figure supplement 3A following prolonged treatment with SKF38393 (25μM) (up to 24h) are shown. **(B)** Original blots of p4E-BP and 4E-BP following SKF38393 (25μM) with/without pretreatment with rapamycin for Figure supplement 3B. (C) Original immunoblots of pAMPK and AMPK for Figure supplement 3C.

**Figure 7 supplement 1.**
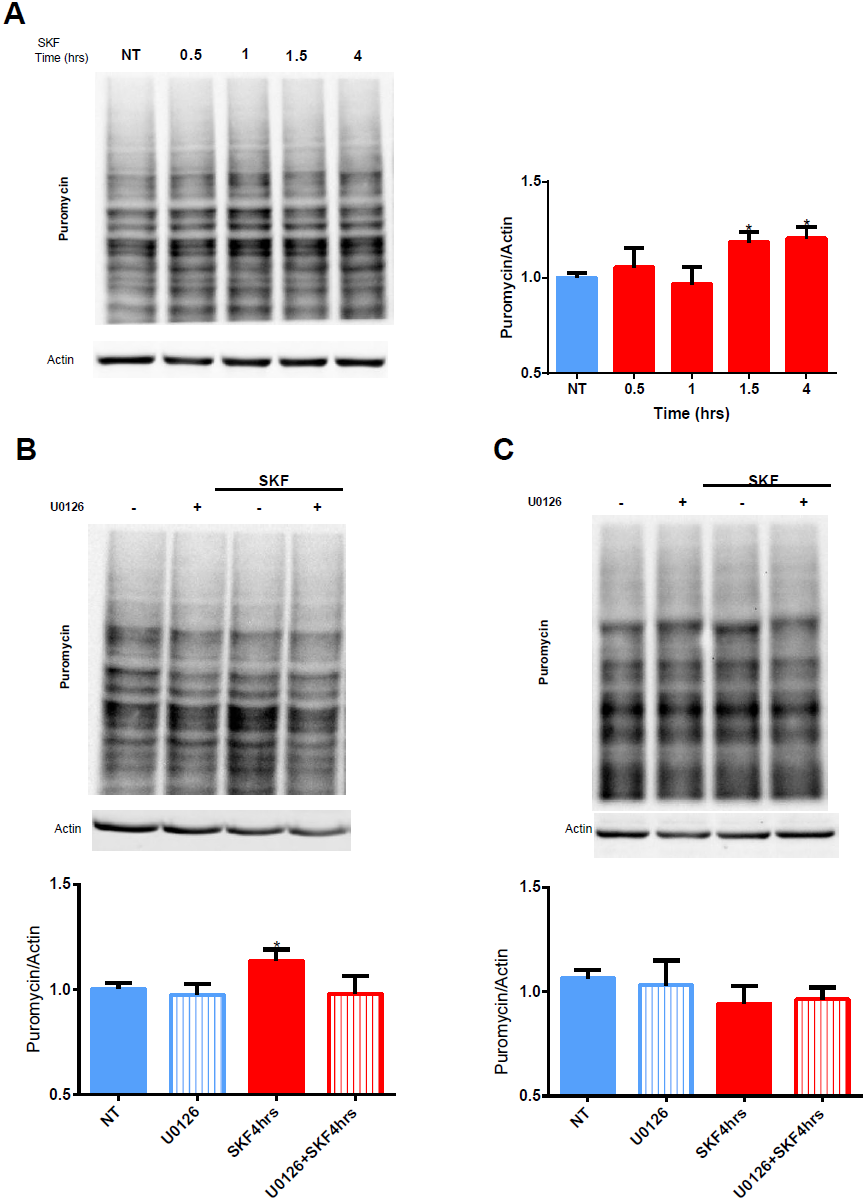
MEK/ERK inhibition reduces D1 receptor-dependent *de novo* protein synthesis increase in WT but not eEF2K-KO mice. **(A)** Time-dependent increase in protein synthesis in primary cultures from C57BL/6 mice treated with SKF38393 (25μM) for the indicated time periods. One way ANOVA F_(4,20_)=3.280, p=0.03. Post-hoc test compared to control; SKF1.5h, p=0.04; SKF4h, p=0.0.04. **(B-C)** WT or eEF2K-KO mouse-derived cortical neurons were pre-treated with MEK inhibitor U0126 (20μM) (or vehicle) for 30 min followed by 4 hours of incubation with SKF38393 (25μM). Cells were treated for 10 min with puromycin (1μg/ml) before harvesting. Puromycin incorporation was detected by Western blotting. Data are means ± SEM of five independent cultures for wild-type and 3 for eEF2K-KO mice. ^*^p<0.05. For WT: One way ANOVA F_(3,44)_=3.288, p=0.02. Post-hoc test compared to Control; SKF4h, p=0.03. For KO: F_(3,25)_=0.614, p=0.61.

**Figure 7 supplement 2.**
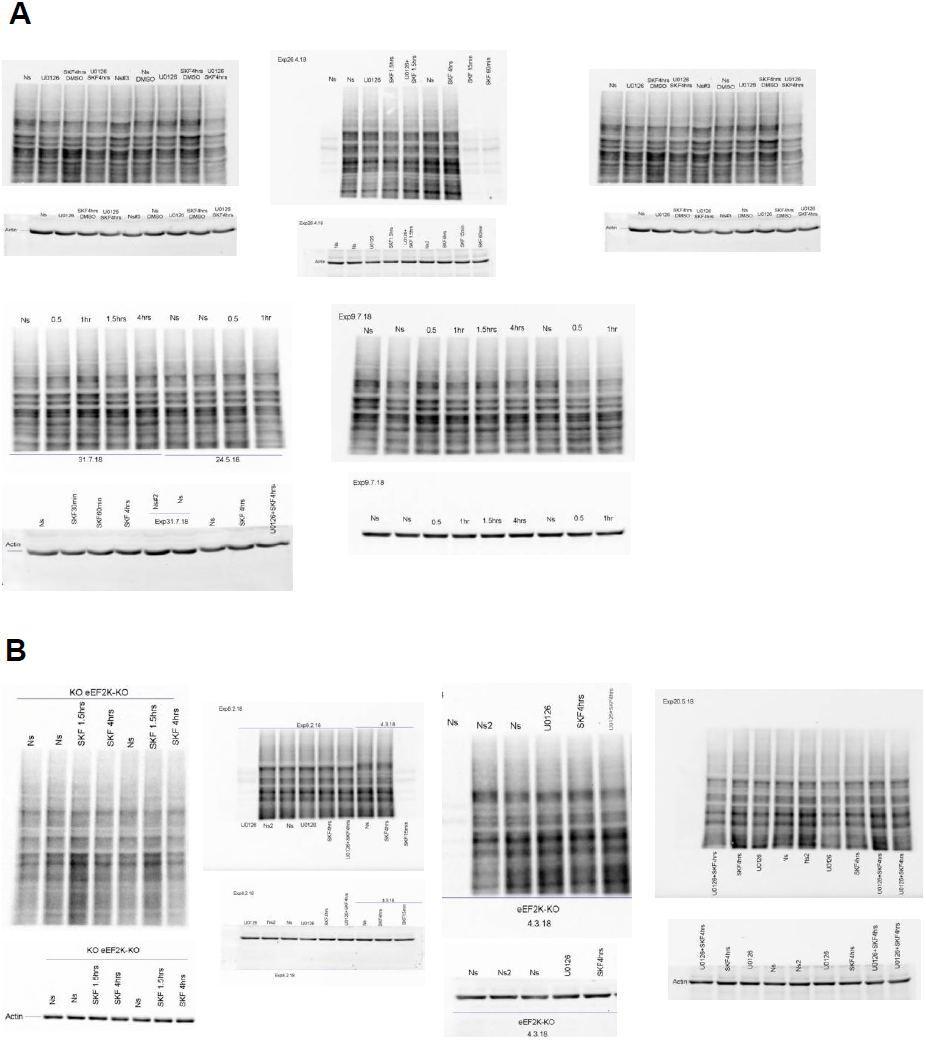
Uncropped original immunoblots of Figure 7 supplement 1. **(A)** Original immunoblots for Figure 7A following prolonged stimulation of SKF38393 (25μM) (up to 4h) followed by incubation with puromycin (1μg/ml) for 10 min. **(B)** Original immunoblots of puromycin from eEF2K-KO mice cultures and their littermates following SKF38393 (25μM) with or without preincubation with MEK inhibitor U0126 (20μM) for 30 min, followed by incubation with puromycin (1μg/ml) for 10 min.

